# Variations in MHC-DRB1 exon2 and associations with Brucellosis susceptibility in Chinese Merino sheep

**DOI:** 10.1101/038539

**Authors:** Yue’e Chen, Wanyun Xu, Chuangfu Chen, Hugh T Blair, Jianfeng Gao

## Abstract

The aim of this study was to investigated the association between MHC-DRB1 exon2 and Brucellosis susceptibility in Chinese Merino sheep. MHC-DRB1 exon2 was amplified by polymerase chain reaction (PCR) from 126 healthy and 67 Brucellosis-infected Chinese Merino sheep. PCR products were analyzed using the SSCP technique, and then cloned to allow sequencing of the different alleles. For each SNP, allelic and genotypic frequencies were compared between case and control samples, in addition the association with Brucellosis susceptibility was determined. Haplotypes and their frequencies were established and analyzed by SHEsis online software. There were forty-one single nucleotide polymorphisms (SNPs) in the 270 bp DNA sequence. The distribution of C>T alleles at locus 109 was significantly different between case and control samples. The linkage disequilibrium (LD) analysis showed that there were nine LD blocks in MHC-DRB1 exon2 and strong LD between SNPs existed in every Block. Haplotype analysis identified nine haplotypes with strong LD, but only Hap8 and Hap9 in case-control groups were significantly different (*P*<0.05); neither haplotype contained the C>T allele at locus 109. In conclusion, genetic variants of MHC-DRB1 gene exon2 demonstrated associations with Brucellosis susceptibility, indicating that further research is warranted.

**Abbreviations:**

MHC: Major Histocompatibility Complex
OLA: Ovine Lymphocyte Antigen
SNP: Single Nucleotide Polymorphism

## Introduction

Brucellosis, caused by the *Brucella* genus, is a widespread, chronic, zoonotic disease of animals and humans (Corbel, 1997). Xinjiang Province in Western China is an important source of production animals, including sheep, cattle, goats and deer. Chinese Merino sheep are one of the most populous breeds in Xinjiang. The presence of *Brucella* is serious, not only because of the harmful effect on human health, but also because infected sheep have lower production. Consequently, there is interest in finding effective means to both prevent *Brucella* infections and to reduce the presence of the *Brucella* organism in Xinjiang. *Brucella* is considered a major health problem requiring urgent action.

The Major Histocompatibility Complex (MHC) is a genetic region comprised of clustered genes which control the regulation of immune response (Ji et al., 2010). Two groups of cell-surface glycoproteins, termed Class I and Class II molecules, are the major MHC-encoded effector molecules and they are intimately involved in T- and B-lymphocyte-mediated immune reactions. The ovine MHC, or Ovine Lymphocyte Antigen (OLA), harbors clusters of immunological genes involved in overall resistance/susceptibility of animals to infectious diseases (Danchin et al., 2004; Flajnik and Kasahara, 2001; Kaufman, 2002). The ovine DRB1 locus is located in the MHC Class II region, and its function is to present extracellular-derived peptides to the immune system. To date, associations between ovine Class II genetic markers and disease susceptibility have been limited to the MHC Class II Orar-DRB1 locus. Reported relationships in sheep include bovine leukemia virus disease, Maedi-Visna and hydatidosis (Larruskain et al., 2010; Li et al., 2011). The majority of nucleotide polymorphisms in Class II loci locate to exon2, consequently, this region has been the principal target in genotyping studies designed to associate MHC genetic diversity with susceptibility to disease (Fallin et al., 2001; Konnai et al., 2003).

Single Nucleotide Polymorphisms (SNPs) belong to the third generation molecular marker technology. Each non-synonymous SNP (nsSNP) changes one amino acid in the gene product causing Single Amino-acid Polymorphisms (SAP) (Schaefer et al., 2012). SNPs are useful genetic markers and can assist in searching for genetic risk factors associated with complex diseases. The application of SNPs in the recognition and identification of disease susceptibility genes has become a key research area. Following completion of the Human Genome Project in 2003 and the International Human Genome Haplotype Map (HapMap) in 2005, the HapMap offered an important additional tool to discover genetic variants associated with diseases. Research using SNPs and haplotypes will play an important role in exploring genetic and pathogenic mechanisms of complex diseases.

Individual SNPs play only a minor role in explaining the genetic variation in complex diseases. However, the genetic information provided by haplotypes is more useful in describing the polygenic nature of genetic diseases (Zaykin et al., 2002). As a consequence, haplotype analysis is increasingly becoming the preferred method for the study of complex diseases (Dukkipati et al., 2006; Sayers et al., 2005). If a specific haplotype can be shown to be distributed differently between case and control samples, this provides strong evidence that the haplotype is associated with the disease, and provides a shortcut for polygenic disease research. While haplotype analysis has been applied to study of the human MHC (Yang et al., 2004; Zhang et al., 2003), this research technique has not been widely applied to investigate the association between animal MHC haplotypes and disease susceptibility.

The purpose of this study was to investigate associations between SNPs and haplotypes of the MHC-DRB1 exon2 in Chinese Merino sheep and Brucellosis susceptibility.

## Results

The Rose Bengal Plate Agglutination Test (RBPT) was used to detect Chinese Merino sheep that expressed antibodies to *Brucella* and were thus considered positive for *Brucella* infection (Figure 1). Out of 193 sheep tested, 126 (65%) were negative for *Brucella*, and 67 (35%) tested positive.

**Figure 1.**
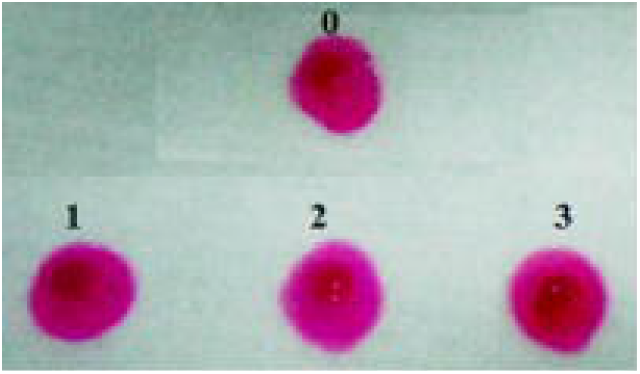
Detection results of Brucella. 1-3: The positive Brucella. 0: Positive control.

PCR amplification products of MHC-DRB1 exon2 were examined via SSCP electrophoresis (Figure 2). Due to the PCR-SSCP method having high sensitivity, many SNP genotypes were detected. Some genotypes were only detected in one individual, and it would be useful to expand the number of sheep sampled to be confident these low frequency SNPs are real.

**Figure 2.**
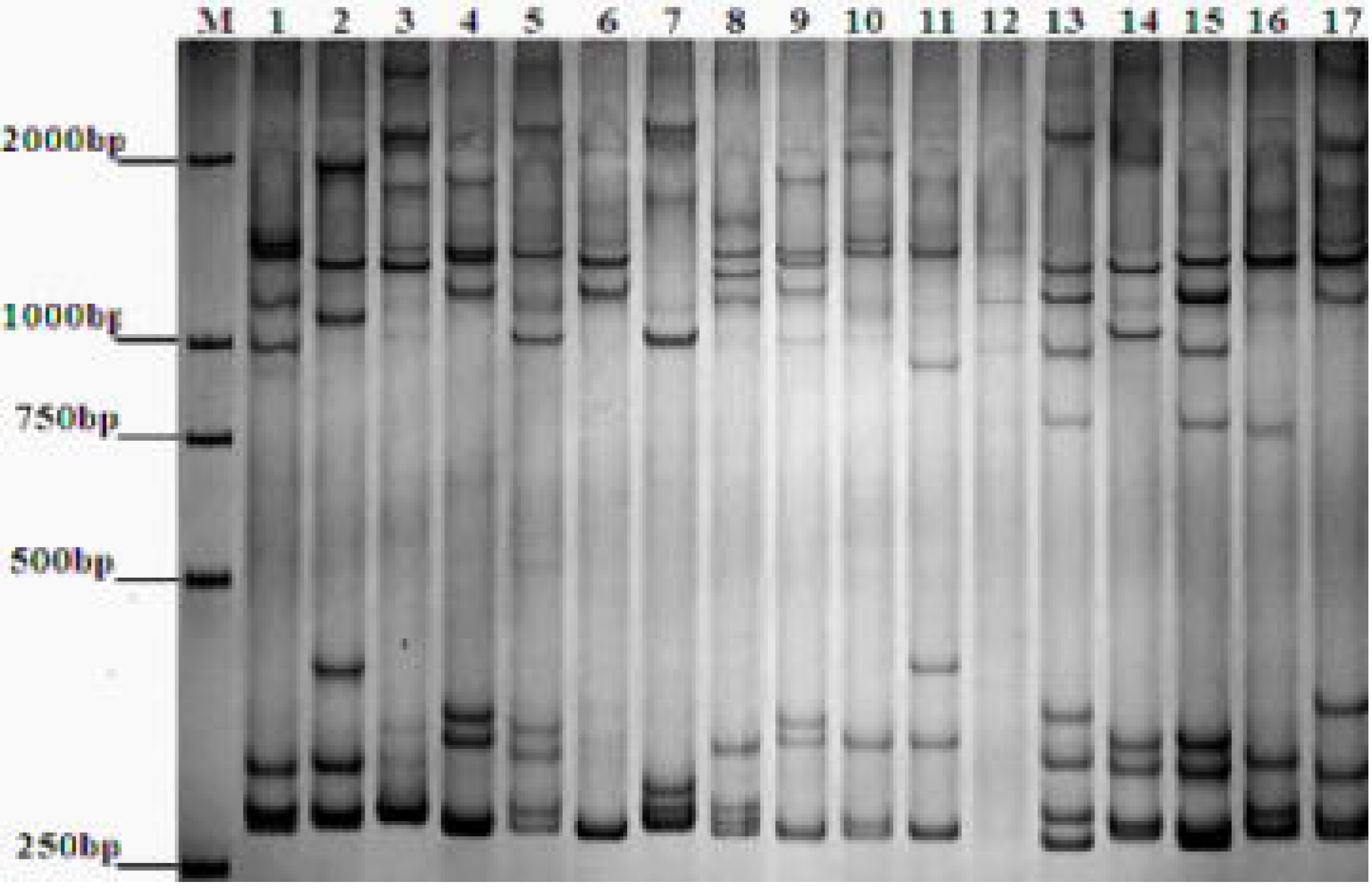
PCR-SSCP detection results of MHC-DRB1 exon2. M: DNA Marker DL2000. 1-17: Different individuals DRB1 exon2 of genotype.

In this study, the PCR-SSCP technique and sequence alignment results demonstrated that MHC-DRB1 gene exon2 of Chinese Merino sheep was richly polymorphic. Sequence alignments were analyzed by GeneDoc software, and the results of alignment are shown in Figure 3. A total of forty-one SNPs were identified in the 270bp length. SNP allelic and genotypic frequencies for case and control groups are shown in Table 1 and Table 2, respectively. The Hardy-Weinberg equilibrium tests showed no significant differences (*P*>0.001).

**Figure 3.**
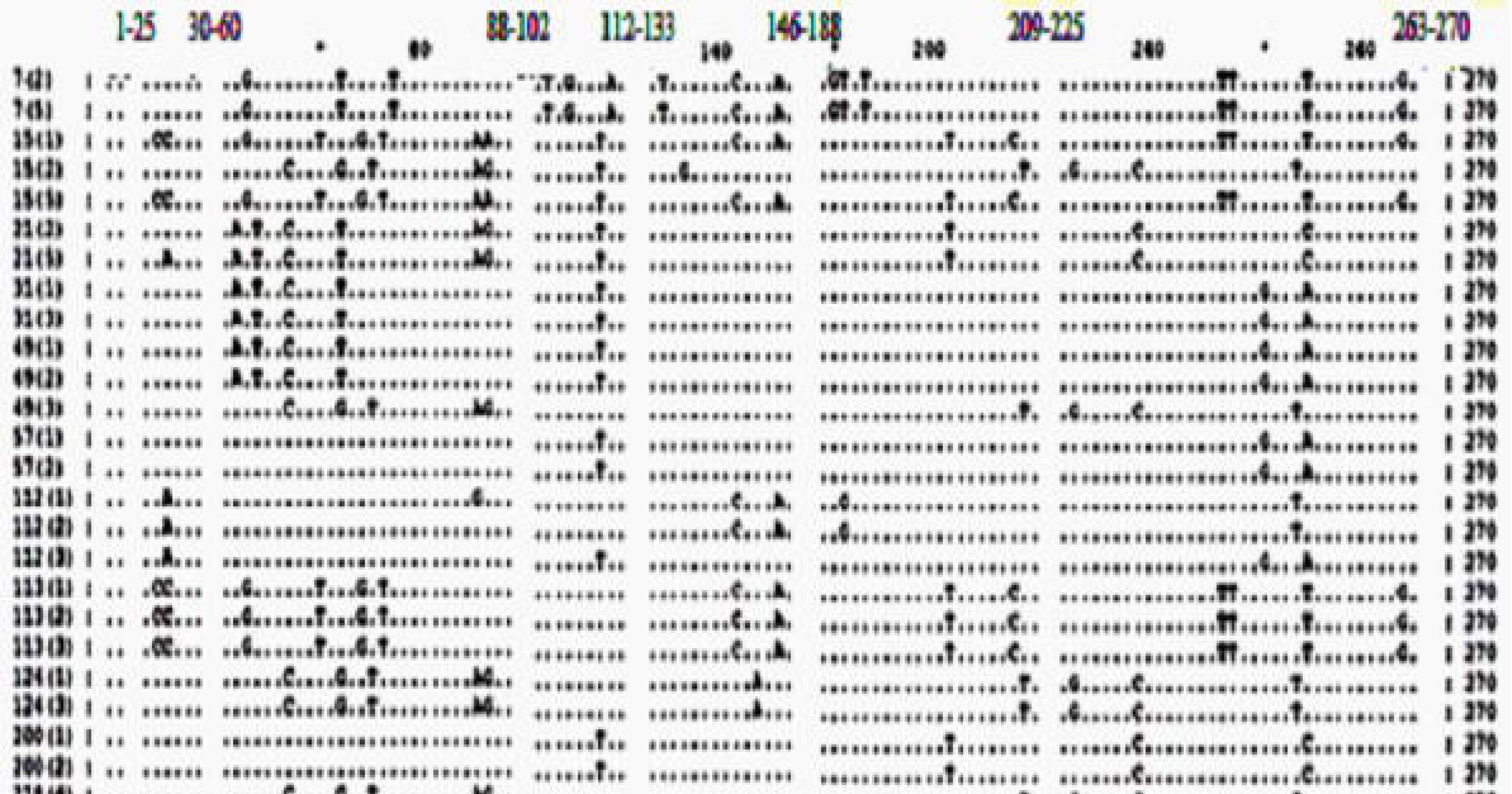
Nucleotide sequence alignment of MHC-DRB1 gene exon2 alleles from Chinese Merino Sheep. Note: The corresponding blank figures represent no polymorphic locus and omit the sequence in the figure.

**Table 1.**
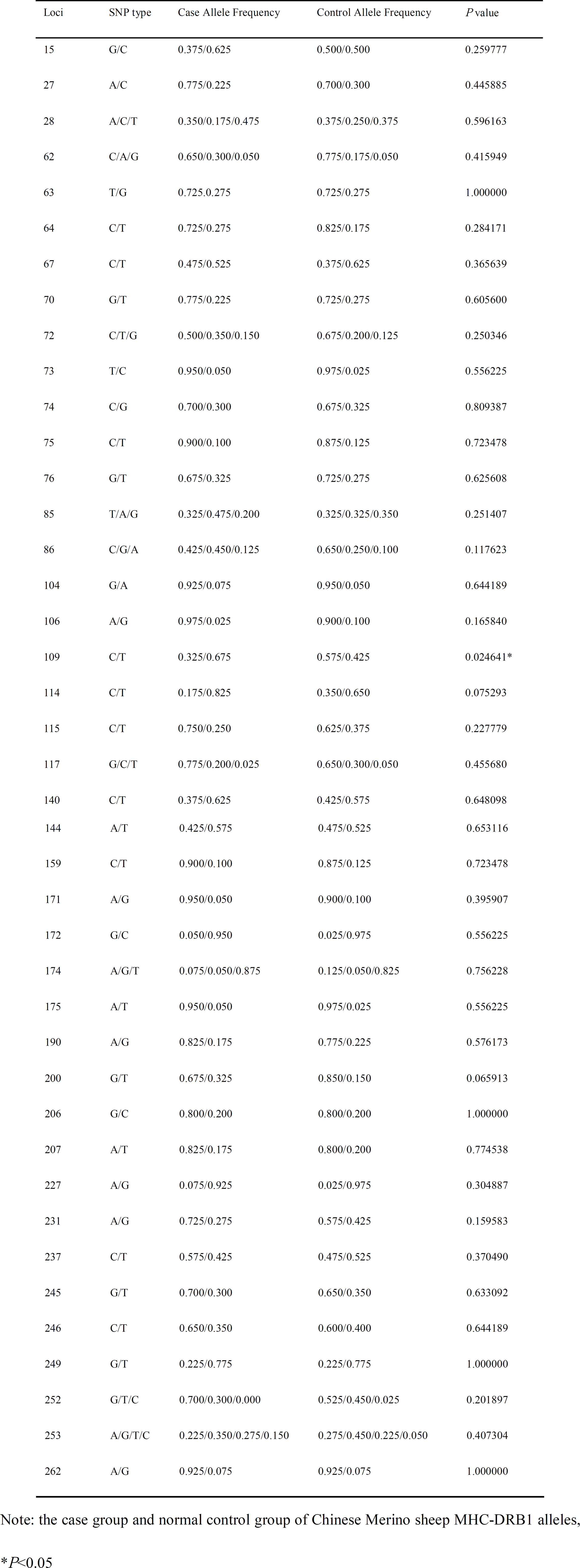
Allele frequencies of PCR-SSCP products about MHC-DRB1 exon2

**Table 2.** Genotype frequencies of PCR-SSCP products about MHC-DRB1 exon2

| Loci | AH | AL | Genotype frequency in case | Genotype frequency in control | *P* value |
| --- | --- | --- | --- | --- | --- |
| G15C | C | G | 0.550/0.150/0.300 | 0.350/0.300/0.350 | 0.374 222 |
| A27C | A | C | 0.700/0.150/0.150 | 0.500/0.400/0.100 | 0.208 108 |
| T63G | T | G | 0.550/0.350/0.100 | 0.450/0.550/0.000 | 0.213 430 |
| C64T | C | T | 0.550/0.350/0.100 | 0.650/0.350/0.000 | 0.338 465 |
| C67T | T | C | 0.300/0.450/0.250 | 0.450/0.350/0.200 | 0.618 440 |
| G70T | G | T | 0.550/0.450/0.000 | 0.450/0.550/0.000 | 0.527 109 |
| T73C | T | C | 0.900/0.100/0.000 | 0.950/0.050/0.000 | 0.548 327 |
| C74G | C | G | 0.500/0.400/0.100 | 0.450/0.450/0.100 | 0.945 797 |
| C75T | C | T | 0.800/0.200/0.000 | 0.800/0.150/0.050 | 0.564 718 |
| G76T | G | T | 0.500/0.350/0.150 | 0.550/0.350/0.100 | 0.883 548 |
| G104A | G | A | 0.850/0.150/0.000 | 0.900/0.100/0.000 | 0.632 607 |
| A106G | A | G | 0.950/0.050/0.000 | 0.850/0.100/0.050 | 0.485 672 |
| C109T | T | C | 0.500/0.350/0.150 | 0.400/0.350/0.250 | 0.139 499 |
| T114C | T | C | 0.700/0.250/0.050 | 0.450/0.150/0.400 | 0.249 167 |
| C115T | C | T | 0.600/0.300/0.100 | 0.450/0.350/0.200 | 0.556 504 |
| C140T | T | C | 0.350/0.550/0.100 | 0.350/0.450/0.200 | 0.648 344 |
| T144A | T | A | 0.350/0.450/0.200 | 0.250/0.550/0.200 | 0.765 928 |
| C159T | C | T | 0.800/0.200/0.000 | 0.750/0.250/0.000 | 0.704 969 |
| A171G | A | G | 0.950/0.000/0.050 | 0.850/0.100/0.050 | 0.347 999 |
| C172G | C | G | 0.900/0.100/0.000 | 0.950/0.050/0.000 | 0.548 327 |
| A175T | A | T | 0.900/0.100/0.000 | 0.950/0.050/0.000 | 0.548 327 |
| A190G | A | G | 0.700/0.250/0.050 | 0.650/0.250/0.100 | 0.830 950 |
| G200T | G | T | 0.550/0.250/0.200 | 0.800/0.100/0.100 | 0.237 128 |
| G206C | G | C | 0.600/0.400/0.000 | 0.650/0.300/0.050 | 0.515 377 |
| A207T | A | T | 0.700/0.250/0.050 | 0.650/0.300/0.050 | 0.938 030 |
| G227A | G | A | 0.900/0.050/0.050 | 0.950/0.050/0.000 | 0.598 389 |
| A231G | A | G | 0.600/0.250/0.150 | 0.400/0.350/0.250 | 0.441 902 |
| C237T | C | T | 0.350/0.450/0.200 | 0.350/0.250/0.400 | 0.289 936 |
| G245T | G | T | 0.600/0.200/0.200 | 0.500/0.300/0.200 | 0.747 584 |
| C246T | C | T | 0.600/0.100/0.300 | 0.450/0.300/0.250 | 0.283 728 |
| T249G | T | G | 0.700/0.150/0.150 | 0.650/0.250/0.100 | 0.691 758 |
| A262G | A | G | 0.900/0.050/0.050 | 0.850/0.150/0.000 | 0.362 661 |

Note: the genotype representation: DD/Dd/dd (D for high frequency allele; D for high frequency allele). AL: low frequency allele name; AH: high frequency allele name

Insertion and deletion sites did not exist in these SNPs. There were nine sites belonging to the PIP and thirty-two belonging to the SP. There were 18 SP conversion sites, including 7 A/G and 11 T/C, and 14 transversion sites, including 1 C/A, 6 G/T, 4 C/G and 3 A/T. The conversion rate was higher than the transversion rate because the cytosine residue CpG dinucleotide is the most frequently mutated sites in the genome. Most of them are methylated, and can spontaneously deaminate resulting in the formation of thymine, which is consistent with the reported results.

The sequence length of MHC-DRB1 exon2 is 270bp, and it can be translated into 89 amino acids. Amino acid sequence analysis showed there were twenty-three mutations causing amino acid changes (Table 3), eight nonsense-mutations and three sense-mutations. For Chinese Merino sheep MHC-DRB1 exon2 of allelic gene sequences of amino acid homology analysis, the homology was more than 81%. The results reflected the consistent with MHC-DRB1 nucleotide polymorphism loci.

**Table 3.** Variable sites and amino acid changes of Chinese Merino sheep MHC-DRB1 exon2

| Number | SNP type | AA change | Number | SNP type | AA change |
| --- | --- | --- | --- | --- | --- |
| 1 | 15G>C | His/Gln | 22 | 140C>T | Leu/Pro |
| 2 | 27A>C | His/Gln | 23 | 144A>T | Gly |
| 3 | 28A>C>T | Arg/- | 24 | 159C>T | Arg |
| 4 | 62C>A>G | Pro/His | 25 | 171A>G | Ala |
| 5 | 63T>G | Pro/His | 26 | 172G>C | Arg |
| 6 | 64C>T | Arg/Trp | 27 | 174A>G>T | Arg |
| 7 | 67C>T | Pro/Ser | 28 | 175A>T | Thr/- |
| 8 | 70G>T | Ala/Ser | 29 | 190A>G | Arg/Gly |
| 9 | 72C>T>G | Ala/Ser | 30 | 200G>T | Cys/Phe |
| 10 | 73T>C | Cys/Ser | 31 | 206G>C | Gly/Ala |
| 11 | 74C>G | Cys/Ser | 32 | 207A>T | Gly/Ala |
| 12 | 75C>T | Cys/Ser | 33 | 227A>G | Arg/His |
| 13 | 76G>T | Ala/Ser | 34 | 231A>G | Tyr/- |
| 14 | 85T>A>G | Ser/Ala | 35 | 237C>T | Asn |
| 15 | 86C>A>G | Ser/Ala | 36 | 245G>T | Leu/Arg |
| 16 | 104G>A | Ser/Asn | 37 | 246C>T | Leu/Arg |
| 17 | 106A>G | Thr/Ala | 38 | 249G>T | Ser |
| 18 | 109C>T | Trp/Arg | 39 | 252G>T>C | Phe/Arg |
| 19 | 114C>T | Arg | 40 | 253A>G>T>C | Asp/Tyr/Asn |
| 20 | 115C>T | Pro/Ser | 41 | 262A>G | Gly/Arg |
| 21 | 117G>C>T | Pro/Ser |  |  |  |

“ - ” : means termination codon

The MHC-DRB1 exon2 sequences of different sheep breeds were aligned through GenDoc software. The results (Table 4) demonstrated that in different studies, the number of SNPs and SAPs were different. However, in the majority of studies between 50% and 70% of SNPs caused amino acid changed. The current study was at the upper limit of this range whereby 30 out of 41 (73%) SNPs resulted in amino acid changes.

**Table 4.** The number of SNP and amino acid mutation loci in different sheep breeds

| Breeds | Authors | Submit date | SNPs No. | SAPs No. | SAPs/SNPs |
| --- | --- | --- | --- | --- | --- |
| Arta | Theodorou,G., Spetsarias,S. | 2012/6/25 | 34 | 21 | 0.618 |
| Bergamasca | Ballingall,K.T., Steele,P.J., | 2010/4/28 | 23 | 16 | 0.696 |
| Blue Faced Leicester | Stear,M.J.,Stirling,D. | 2008/9/16 | 25 | 17 | 0.680 |
| Kozani | Theodorou,G., Spetsarias,S. | 2012/6/25 | 17 | 12 | 0.706 |
| Lori Bakhtiari | Nikbakht Brujeni,G.,Rezaii,H. | 2010/9/2 | 25 | 15 | 0.600 |
| Prealpe | Ballingall,K.T., Tassi,R., | 2009/5/12 | 34 | 19 | 0.559 |
| Red Maasai | Ballingall,K.T., Steele,P. | 2010/8/25 | 43 | 26 | 0.605 |
| Scottish Blackface | Stirling,D., Stear,M.J. | 2008/9/16 | 58 | 31 | 0.534 |
| Sopravvisana | Ballingall,K.T., Steele,P.J., | 2010/4/28 | 19 | 10 | 0.526 |
| Zandi | Nikbakht Brujeni,G., Rezaii,H. | 2010/9/2 | 19 | 7 | 0.368 |

In this study, the polymorphism of MHC-DRB1 gene exon2 associated with Brucellosis susceptibility, the result of analysis found that MHC-DRB1 was likely to be one of the genes associated with Brucellosis susceptibility, C>T alleles at the 109 locus in the case-control samples distribution existed significant difference (*P*<0.05), and preliminary analysis suggested that MHC-DRB1 exon2 109 C>T associated with Brucellosis susceptibility. Association analysis was conducted for each genotype of the gene polymorphisms, showing that the site of DD/Dd/dd genotype in the case-control samples distribution had no significant difference (*P*>0.05).

Haplotypes generally had more information content than individual SNPs. Therefore, we performed linkage disequilibrium(LD) and haplotype analysis for the SNPs with MAFs<5% and the genotype distributions were in Hardy-Weinberg equilibrium (*P*<0.001) in Chinese Merino sheep. The standardized measure of LD denoted as D’ were calculated for all pairs of SNPs. Among forty-one SNPs in MHC-DRB1 exon2, only twenty-nine SNPs were eligible, and were used to analyze LD in both case and control, the haplotypes LD map as shown in Figure 4, indicated the LD of the SNPs, the number in box such as 99 is 0.99, the greater D’ value means the stronger LD degree between each two SNPs, D’>0.9 is usually considered highly LD, and D’>0.7 is deemed to the two SNPs located in same block, which can perform haplotypes analysis. The block diagram of color from light to deep (white and red), said the LD degree from low to high, deep red means completely linkage. LD analysis found that the MHC-DRB1 exon2 had nine LD Blocks (Table 5), and each two SNPs had strong LD in every Block.

**Figure 4.**
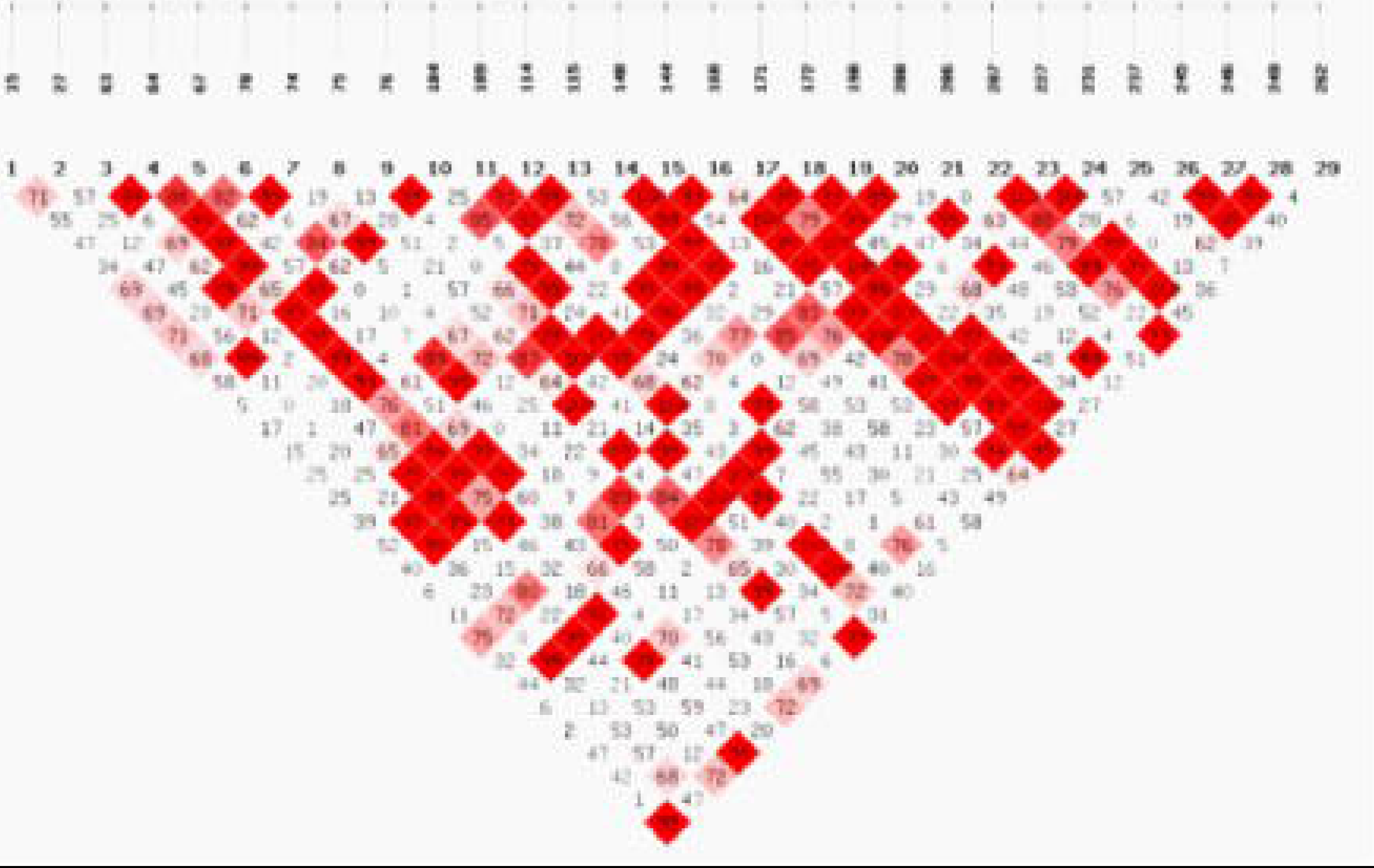
The figure of linkage disequilibrium Haplotypes based case-control sample. Note: The first line number indicates the location of SNPs; and the second line of figures indicate the number of SNPs, a total of 29. The number in graph box is the value of linkage disequilibrium(D’), D’ values greater the darker, which means that the higher the degree of linkage disequilibrium. Conversely represents the lower the degree of linkage disequilibrium.

**Table 5.** The composition of every LD Block with different SNPs

| LD Block | SNPs |
| --- | --- |
| Block1 | T63G、C64T |
| Block2 | C64T、C67T、G70T |
| Block3 | G70T、C74G |
| Block4 | G76T、G104A |
| Block5 | C109T、C114T、C115T |
| Block6 | C140T、A144T、C159T |
| Block7 | A171G、G172C、A190G、G200T |
| Block8 | A207T、A227G、A231G |
| Block9 | G245T、C246T、G249T |

Due to a series of genetic diseases are often not caused by a single SNP loci, but by the combination of SNPs on several sites, therefore, in the study of disease association analysis, based on the multiple sites of SNPs studies tend to have more strength and more convincing than a single SNP loci. Because haplotype frequency cannot less than 0.03 in a population, the haplotype combination which frequency yet reached 0.03 should be ignored during statistics, so that only nine haplotype combinations were analyzed in the MHC-DRB1 exon2 (Table 6), the result of analysis found that Hap8 and Hap9 these two haplotypes frequency in the case group was 12.5% and 15% respectively, was significantly higher than control group, haplotype frequency distribution difference was statistically significant (*P*<0.05), initially speculated that these two haplotypes may associated with Brucellosis susceptibility.

**Table 6.**
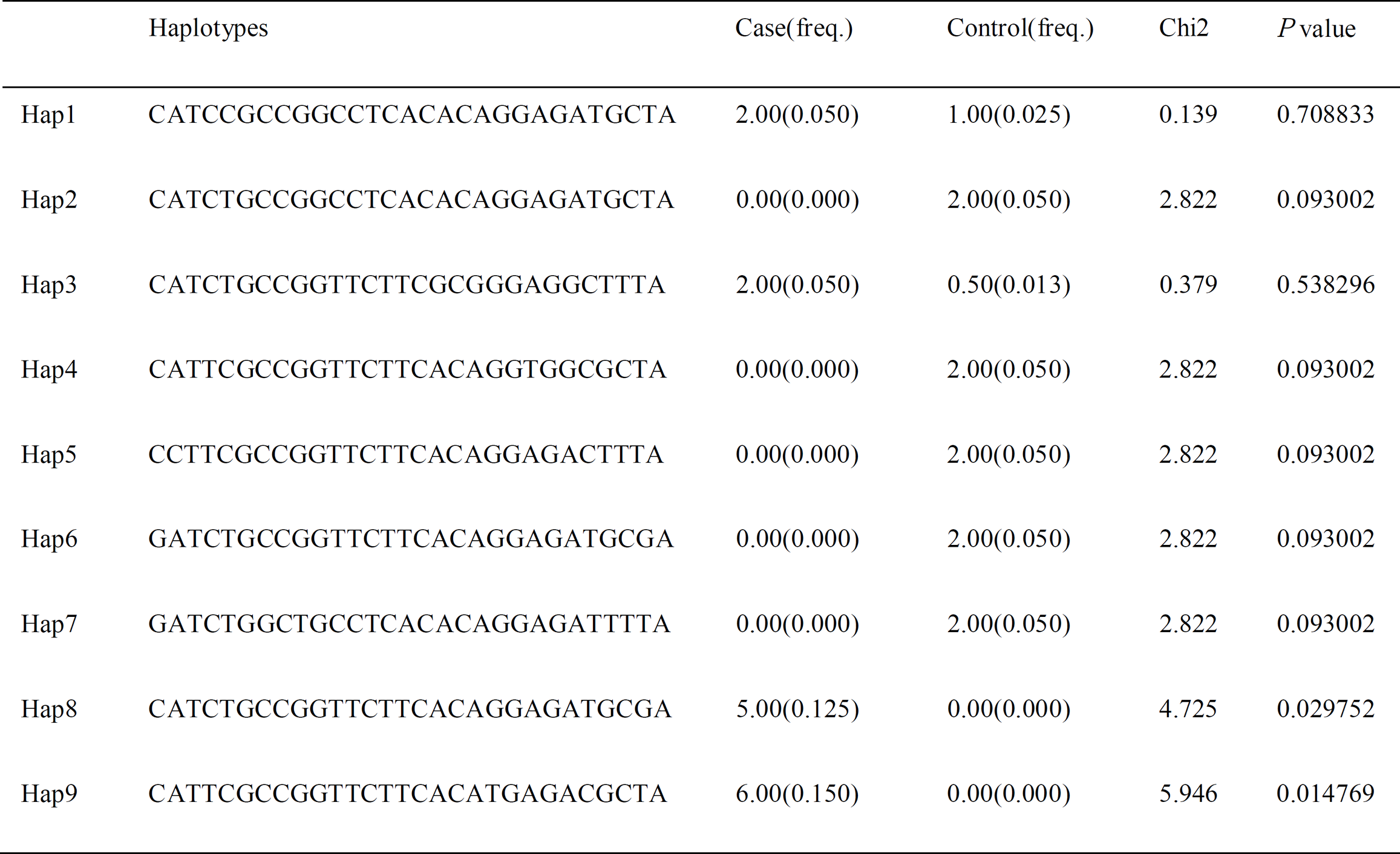
The Haplotypes frequencies of SNPs in case-control MHC-DRB1 exon2

## Discussion

A high degree of polymorphism and base mutation are a very prominent feature of vertebrate MHC genes, having become a hot research topic in livestock resistance breeding. Forty-one SNPs in MHC-DRB1 gene exon2 of Chinese Merino sheep observed in this study are newly identified and have not been reported previously. These results confirmed that PCR-SSCP is a useful tool for easy and efficient identification of DNA polymorphisms and can be employed for evaluating genetic variability in large livestock populations. Furthermore, according to the previously study about SNPs and SAPs, the results fully demonstrated that MHC-DRB1 gene of Chinese Merino sheep is really highly polymorphic.

Compared with other varieties sheep polymorphic in MHC-DRB1 gene exon2, there were a lot of differences in different breeds of sheep. The discrepancy in different populations was probably caused by the following reasons: one the one hand, some polymorphisms can only be preserved and inherited in ancient and special animals. On the other hand, different selection purpose and selection history lead to the diversity.

The main function of MHC is antigen presentation, playing a very important role in the immune system of animal body, and it has a close relationship with the livestock disease susceptibility. Many disease susceptibility are associated with MHC, as a candidate gene for disease susceptibility has become one of the research hotspots in modern molecular immune genetic. Through the study of the association between SNPs and the Brucellosis susceptibility, the 109 C>T was found to be significantly associated with Brucellosis susceptibility, which provide scientific basis for choice Brucellosis susceptibility molecular marker-assisted, genetic loci for further search with susceptibility and lay the foundation for genetic breeding, but about whether the 109 C>T as genetic markers of Brucellosis, also need to attack toxic experiment will be carried out to further verify. After further verification, this SNP could be a useful molecular marker for use in poultry breeding.

In case and control association studies, when LD exists in each two SNP, haplotypes generally had more information content than individual SNPs. This study calculated the degree of LD and inferred haplotype using SHEsis online software. Because of LD of SNPs in very close genetic distance, but in order to successfully identify may lead to changes in the disease, many relational analysis method using multiple high density SNPs sites to study(Epstein and Satten, 2003), on the other hand, haplotype is just a collection of closely linked SNPs on the same chromosome allele, it contains a number of pairwise LD information, so the gene location problem, based on haplotype association are more powerful than based on single SNP loci analysis (Zaykin et al., 2002).

The haplotype analysis found that nine haplotypes were constructed by using twenty-nine SNPs, if LD did not exist between each two SNPs, twenty-nine SNPs should produce 2^29^ haplotypes, but only nine haplotypes were detected among these SNPs in this study because of strong LD. In addition, we also investigated haplotype frequencies between case and control in Chinese Merino sheep, and speculated that Hap8 and Hap9 may associated with the Brucellosis susceptibility which may relate to immune traits association studies and lay the foundation for molecular marker assisted selection. Further studies as well as functional analysis are required to fully elucidate how these interesting gene polymorphisms may affect MHC-DRB1 gene exon2 activity and/or probably act as candidate markers associated with Brucellosis susceptibility in sheep.

## Materials and Methods

### Sample collection and DNA isolation

Blood samples were obtained from 193 Chinese Merino sheep farmed by the Xinjiang Production and Construction Corps Agricultural 9^th^ Division 170 Regiment. Genomic DNA was isolated from whole blood samples with the use of phenol chloroform extraction, and stored at −20°C for later use.

### Detection of *Brucella*

The Rose Bengal Plate Agglutination Test (RBPT) was used to detect *Brucella* antibodies. A serum sample of 30μl was evenly mixed with 30 μl antigen at room temperature, and reaction results were recorded after 4 to 10 min, results were immediately compared to positive serum. Serum agglutination with any degree of granularity or flocculation was designated as positive, while no agglutination was designated as negative.

### Amplification of OLA-DRB1 exon2

Exon2 of OLA-DRB1 (Accession Number: FR848372) was amplified using PCR with the following oligo nucleotide primers: DRB1-1: 5’ TAT CCC GTC TCT GCA GCA CAT TTC 3’ and DRB1-2: 5’ CTC GCC GCT GCA CAC TGA AAC TCT 3’. PCR was performed in a reaction volume of 20 μl containing 50 ng/μl ± 4 ng/ul genomic DNA, 10 μl 2×PCR Master Mix, 1 μl of each primer (10 umol/L) and 7 μl ddH_2_O. The PCR was undertaken in a Mastercycler gradient thermocycler (Eppendorf China Limited) under the following conditions: denaturation at 94°C for 5 min, followed by 94°C for 30 s, 63°C for 1 min, and 72°C for 1 min for 30 cycles, and a final extension at 72°C for 10 min. The amplified fragment of 286 bp consisted of 16 bp of intron1 and the entire exon2 of 270 bp. An aliquot of 5 μl reaction product was used to check the concentration and quality of the PCR products by agarose gel electrophoresis.

### SSCP (Single-strand conformational polymorphism) Analysis

Each PCR product (4 μl) was mixed well with 9 μl of denaturing solution (95% formamide, 20 mM EDTA, 0.05% bromophenol blue and 0.05% xylene-cyanole). Mixtures were submitted to a denaturation step at 95°C for 5 min, and then rapidly chilled on ice. Ten μl of the denatured mixture was directly electrophoresed on precast 10% (Acr : Bis=39:1) non-denaturing polyacrylamide gels for SSCP. After electrophoresis, one glass plate was removed and the gel on the second glass plate was stained in 0.1% AgNO_3_ for 8 min, and then briefly rinsed with distilled water two times, for 20 s each time. Gels were developed in 400 ml containing 1.2% (w/v) NaOH and 0.4% formaldehyde. When the desired band intensity was achieved, development was stopped. Stained gels were rinsed with distilled water and gels were scanned using an X-ray lamp.

### Cloning and sequencing

Following PCR-SSCP analysis, PCR amplification products were recycled and purified by an agarose gel purification kit to enable identification of different genotypes for each individual. Recycled DNA fragments were attached to the pMD19-T vector, and transformed into TOP10 E.coli competent cells on LB plates coated with Amp. Clones were inoculated onto the culture medium containing Amp and allowed to culture. Thus, the bacteria were used as a template for PCR amplification to identify recombinant clones. If they were amplified completely consistently with the objective fragment, they were identified as positive clones. Bacteria containing positive clones were sent to BGI, Beijing, for sequencing. Sequence results were analyzed using DNASTAR and GeneDoc software.

### Data processing and statistical analysis

The genotypic frequency of SNPs from the MHC-DRB1 gene exon2 were analyzed for Hardy-Weinberg equilibrium (HWE) in case and control Chinese Merino sheep using chi-squared with a significance level of *P*=0.001. Comparisons of allelic and genotypic frequencies in case-control groups and association analysis between the polymorphisms and Brucellosis susceptibility were undertaken using SHEsis software.

### Linkage disequilibrium and Haplotype analysis

SNPs whose minor allele frequencies (MAFs) were <5%, failed genotyping due to technical errors, or failed to meet Hardy-Weinberg equilibrium (*P*<0.001) were removed from the haplotype structure. The pattern of Linkage Disequilibrium (LD) between the SNPs was measured by the LD coefficient D’=D/Dmax., The magnitude of LD between matching sites was used to indicate whether or not LD existed. The value of D’ was calculated by the online genetics software SHEsis. This software tests the HWE of the sample population, analyses LD of large samples while testing for multiple SNP loci at the same time and constructs haplotype associations (Li et al., 2009; Yong and Lin, 2005).

## Acknowledgements

We thank everyone from the Xinjiang Production and Construction Corps Agricultural 9th Division, 170 Regiment, who provided samples and their participation and support for this work. We are also warmly appreciative to Ma Runlin Professor in the Institute of Genetics and Developmental Biology, Chinese Academy of Sciences, for his helpful advice and thank BGI, Beijing, for providing the sequencing platform.

